# Primate-specific *cis*- and *trans*-regulators shape transcriptional networks during human development

**DOI:** 10.1101/2021.08.18.456764

**Authors:** Julien Pontis, Cyril Pulver, Evarist Planet, Delphine Grun, Sandra Offner, Julien Duc, Andrea Manfrin, Matthias Lutolf, Didier Trono

## Abstract

The human genome contains more than 4.5 million inserts derived from transposable elements (TE), the result of recurrent waves of invasion and internal propagation throughout evolution. For new TE copies to be inherited, they must become integrated in the genome of the germline or preimplantation embryo, which requires that their source TE be expressed at these stages. Accordingly, many TEs harbor DNA binding sites for the pluripotency factors OCT4, NANOG, SOX2, KLFs and are transiently expressed during embryonic genome activation. Here, we describe how many primate-restricted TEs have additional binding sites for lineage-specific transcription factors driving their expression during human gastrulation and later steps of fetal development. These TE integrants serve as lineage-specific enhancers fostering the transcription, amongst other targets, of KRAB-zinc finger proteins of similar evolutionary age, which in turn corral the activity of TE-embedded regulatory sequences in an equally lineage-restricted fashion. Thus, TEs and their KZFP controllers play broad roles in shaping transcriptional networks during early human development.

## Introduction

The human genome hosts some 4.5 million sequence inserts readily recognizable as derived from transposable elements (TEs), most of them retroelements that replicate through a copy-and-paste mechanism based on the reverse transcription of an RNA intermediate followed by insertion of its DNA copy, whether ERVs (endogenous retroviruses), LINEs (long interspersed nuclear elements), SINEs (short interspersed nuclear elements, which include primate-specific Alu repeats) or SVAs (SINE-VNTR-Alu, composites of an ERV and Alus, restricted to hominids). TEs are increasingly recognized as a major driver of genome evolution owing notably to their recombinogenic and regulatory potential, even though most are unable to spread further due to inactivating mutations^1,2^.

TEs are tightly controlled by epigenetic silencing mechanisms, and as a corollary many are expressed when these mechanisms are put on hold in the context of genome reprograming. Correspondingly, thousands of TEs display marks of open chromatin and are transcribed during embryonic genome activation (EGA)^3-6^, albeit mostly brlongoing to the groups of primate-restricted ERVs (HERV9, HERVK, HERVL, HERVH), SVAs, Alus, and young LINE-1s. The same subset of TEs is enriched in acetylated histone, a hallmark of enhancers, in human embryonic stem cells (hESCs) derived from the pre-implantation embryo^7-9^, where many are bound and controlled by pluripotency transcription factors (TFs)^7,9-13^. This correlates with the TE transcription-dependence of transposition events that must occur in the germline or in preimplantation embryos to be inherited. In return, the broad induction of transcriptionally active TE loci during EGA likely contributes to the efficiency of this process and their *cis*-regulatory influences shapes the gene regulatory landscape of the pre-implantation embryo. In a remarkable regulatory feedback loop, the pluripotency factor-mediated activation of primate-restricted TE-embedded enhancers leads to the expression of evolutionary contemporaneous Krüppel-associated box (KRAB)-containing zinc finger proteins (KZFPs), which act as sequence-specific repressors of these EGA-induced TEs^9,14^.

While KZFP-induced heterochromatin formation and DNA methylation are believed to maintain TE-embedded regulatory sequences in a repressed state at later stages of development and in adult tissues^15-18^, the present study reveals that many of these sequences are expressed during gastrulation and are accessible in fetal tissues, displaying in these settings a high degree of lineage specificity reflecting their activation no longer by pluripotency factors but by cell type-specific transcription factors (TFs). Furthermore, we determine that many of these TE-derived enhancers are enriched near KZFP genes, the secondary stimulation of which is responsible for lineage-restricted heterochromatin formation during germ layer formation. Thus, evolutionary recent TEs and their KZFP controllers strongly influence and thereby confer a high degree of species specificity to the conduct of human gastrulation and fetal development.

## RESULTS

### Cell-type specific expression of primate-restricted TEs during human gastrulation

To examine the transcriptional state of TEs in the immediate post-implantation period, we analyzed single-cell transcriptome data from a gastrulating human embryo^19^. Gene expression patterns allowed the grouping of cells in clusters corresponding to epiblast, primitive streak, primordial germ cells, ectoderm, nascent, emergent, advanced, yolk sac and axial mesoderm, endoderm and early hematopoietic compartment (Fig. 1a, Fig. S1a). Overall, we could detect transcripts emanating from more than 100,000 TE loci, with a marked relative overrepresentation of primate-specific integrants (Fig. 1b, Fig. S1b). Many of these evolutionary recent TEs belong to the SVA, HERVK, HERVH and L1PA3, L1PA2 and L1Hs subgroups (Fig 1c), previously found to be transcribed during embryonic genome activation (EGA)^3,9^. However, these EGA-induced subfamilies were more highly expressed in the primitive streak compared to the epiblast, consistent with *de novo* transcription, and some of them further displayed cell-specific patterns of expression (Fig 1d and Fig S1c-d). For instance, HERVH expression was broad but highest in PGC and axial mesoderm and low in hematogenic derivatives, whereas HERVK transcripts were most abundant in PGCs, endodermal cells, and blood progenitors. Interestingly the evolutionary young LTR5Hs-derived HERVK were expressed in both PGCs and endodermal cells while the more ancient LTR5B-derived HERVK were detected only in the latter tissue (Fig 1d, Fig S1c-d). Moreover, HERVK11 and HERVS71/HERVK22 transcripts, not previously detected during EGA, were also not present in epiblast cells but were abundant in the primitive streak and nascent mesoderm for the former and in definitive endoderm for the latter, and HERVIP10FH and HERV17 integrants were similarly *de novo* expressed in PGCs (Fig 1d, S1cd).

**Figure 1.**
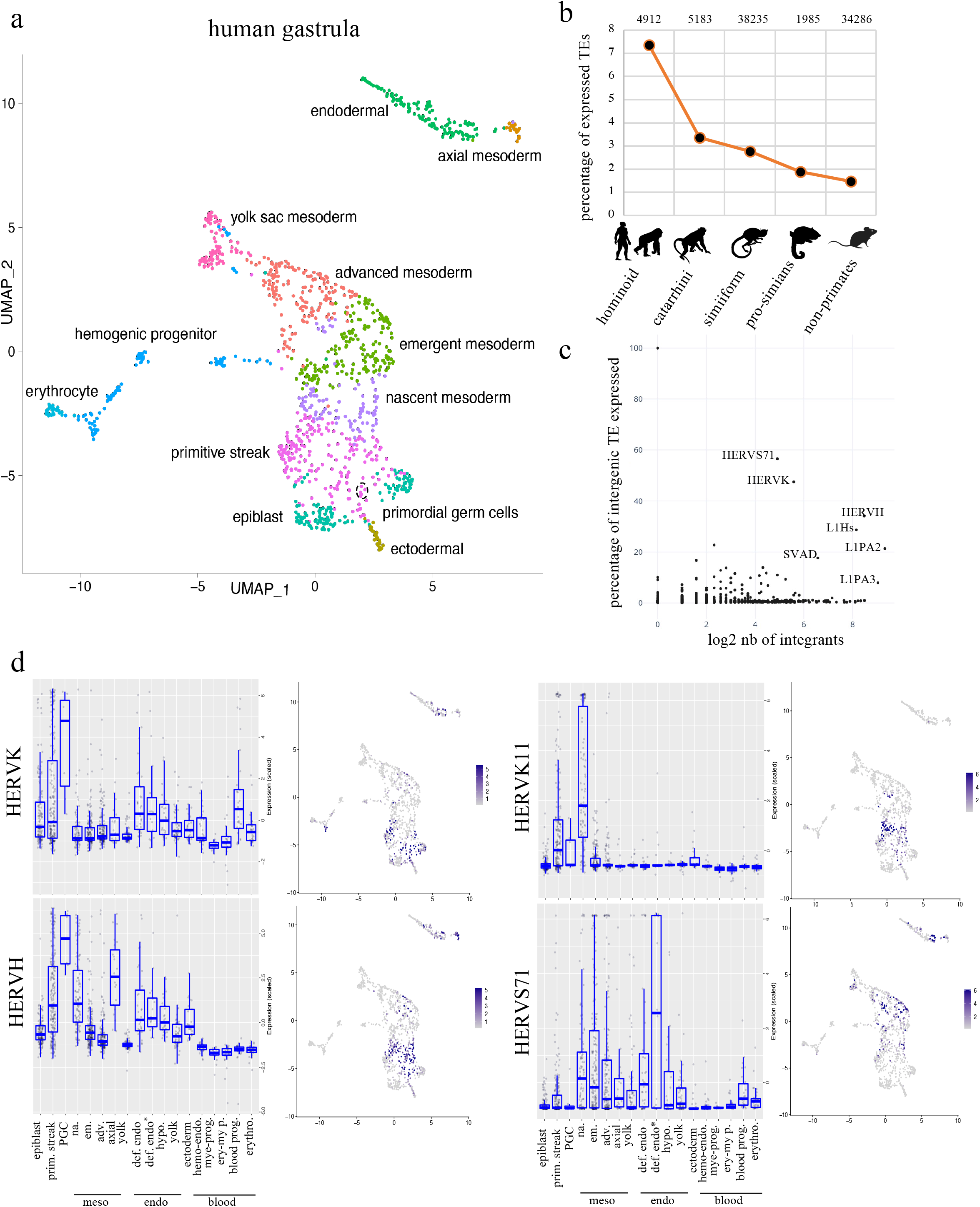
Cell-type specific expression of primate-restricted TEs during human gastrulation. **a**, Cellular composition of the human gastrula. UMAP (Uniform Manifold Approximation and Projection) based on gene expression in single cells from human gastrula; colors correspond to the different cell types identified in^19^. **b**, Age distribution of TEs expressed in human gastrula. All examined TE subfamilies (excluding DNA transposons) were assigned their evolutionary age category, and percentage of expressed integrants from each age category was plotted. On top are plotted the number of expressed integrant. **c**, Relative expression of TE subfamilies in human embryo, depicting number (x-axis) and percentage (y-axis) of integrants expressed from indicated subfamilies. **d**, Cell-specific expression of indicated TE subfamilies; each dot represents normalized expression of TE in one cell, grouped in boxplots corresponding to one specific cell type of human gastrula sub-clustering: epiblast, primitive streak (prim. streak), primordial germ cells (PGC), nascent mesoderm (na.), emergent mesoderm (em.), advance mesoderm (adv.), axial mesoderm (axial), yolk mesoderm (yolk), definitive endoderm (def. endo.), definitive endoderm non-proliferative (def. endo*), hypoblast (hypo), yolk endoderm (yolk), ectoderm (ectoderm), hemogenic endothelium (hemo-endo.), myeloid progenitor (mye-prog.), erythro-myeloid progenitor (ery-my p.), blood progenitor (blood prog.), erythrocyte (erythro.); UMAPs represent the relative TE subfamily expression distribution in the human gastrula clustered on gene expression.

### Evolutionary recent TEs maintain their cis-regulatory potential during human gastrulation and fetal development

Having documented the germ layer-specific expression of EGA-induced and several other TE subfamilies during human gastrulation, we analyzed a time-course single-cell transcriptomic of hESC-derived embryoid body^20^. The results confirmed the lineage-specific expression of several TE subfamilies in this *in vitro* system (Fig 2a, S2a). Moreover, upon examining chromatin in purified hESC-derived PGC-like and endodermal cells, we observed cell-type-specific accessibility patterns that correlated with expression for specific gastrula TE subfamilies (Fig 2b).

**Figure 2.**
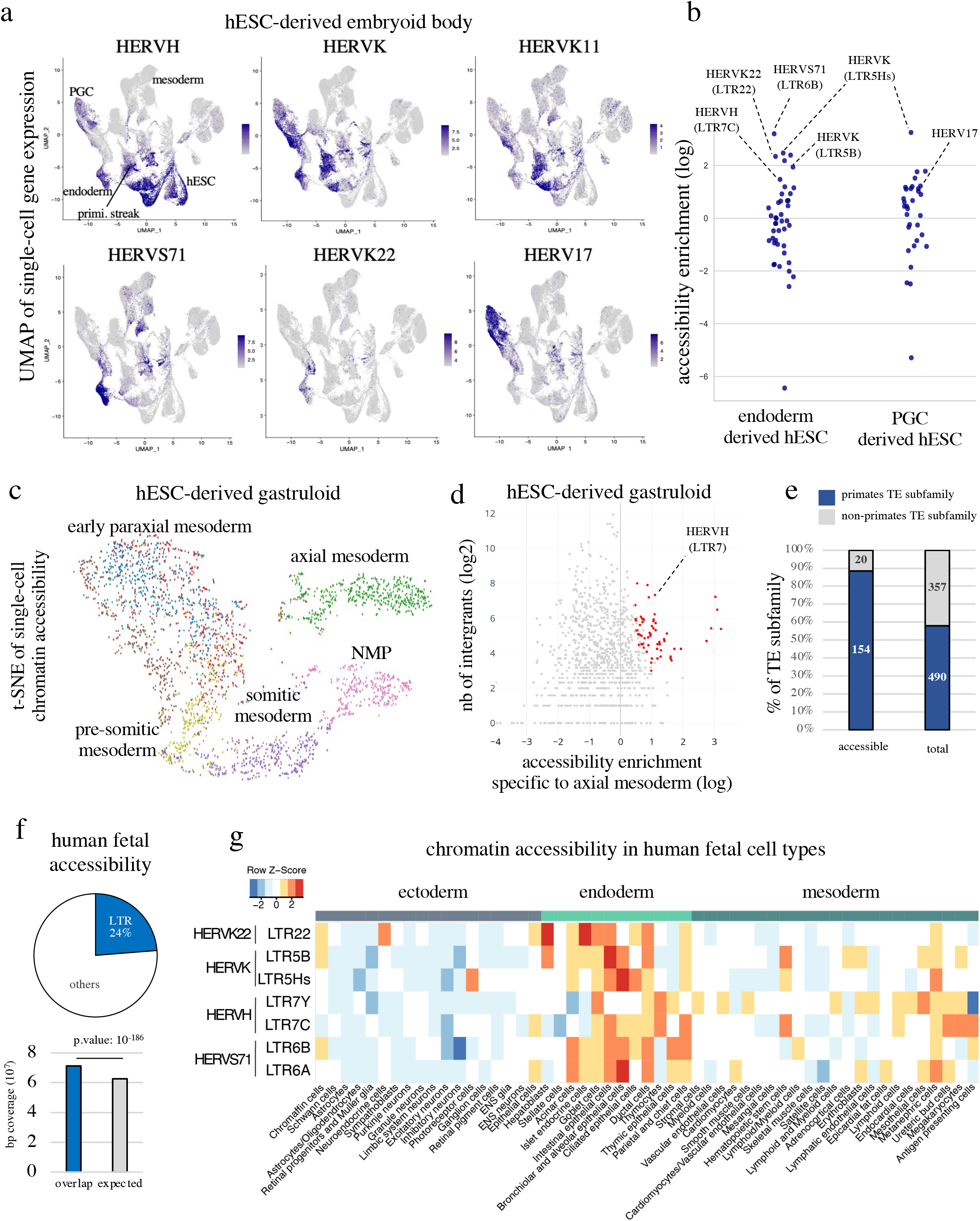
Evolutionary recent TEs maintain their cis-regulatory potential during human gastrulation and fetal development. **a**, Cell-type-specific expression of TE subfamily during embryoid body differentiation. Each plot represents a UMAP of single-cell gene expression during *in vitro* differentiation of hESC into embryoid body over 5 days re-analyzed from ^20^; hESC corresponds to day 0; primitive streak (prim. streak) to days 1-2 with TBXT expression; cells at days 2-5 are stratified in PGC expressing NANOS3, NANOG, and SOX17, endoderm expressing SOX17 and GATA6, and mesoderm expressing GATA6 only. Color scale corresponds to level of relative TE subfamily expression based on normalized read counts. **b**, Chromatin accessibility at endodermal and PGC expressed TE subfamilies (adjusted p-value < 0.05) in hESC-derived endoderm and PGC, calculated over random genomic distribution and represented in natural log. **c**, Chromatin accessibility in human gastruloids, analyzed at the single-cell level. t-SNE plot (t-distributed Stochastic Neighbor Embedding) represent chromatin accessibility clustering and colors correspond to gene expression clustering from S2c-e. **d**, TE subfamily enrichment of chromatin accessibility (p-value < 0.05) in axial mesoderm. Each dot represents a TE subfamily; x-axis, natural log fold enrichment TE loci compared to a random genomic distribution and y-axis represent the log2 number of accessible integrants. Red dots are TE subfamilies with a p-value of enrichment < 0.05. **e**, Relative accessibility of primate or non-primate TE integrants in human pre-implantation embryo, fetus and *in vitro* hESC differentiation model re-analyzed from^4,22,23^, indicating number of TE subfamilies with significant accessibility in at least one developmental context (50 accessible integrants with > 2 fold enrichment over random genome coverage, p-value < 0.05). **f**, Contribution of LTR TEs to chromatin accessibility at fetal stage. Top, distribution of all chromatin accessible sites in human fetus (more than 1 million loci) and the proportion overlapping LTR; bottom, measured vs. expected contribution of LTR-derived TE. **g**, Enrichment over random distribution of selected TE subfamilies in indicated cell types during fetal development re-analyzed from^22^; unknown, maternal and placental cell types were removed.

To investigate further the link between TE subfamily expression and chromatin accessibility changes during gastrulation, we generated gastruloids, an elongated self-organizing embryoid body recently proposed as model for studying some aspects of human gastrulation^21^. Using multi-omics assay to identify both chromatin accessibility and RNA content at the single nucleus scale, we could identify five main tissues in this experimental system: TBXT-expressing axial mesoderm, SOX2-expressing neuro-mesenchymal progenitors (NMP), and three different stages of paraxial mesoderm differentiation preferentially expressing respectively TBX6 (somitic), or PAX3/TWIST1/SIX1 (early paraxial and pre-somitic) with different degree of somitic HOX gene markers (Fig 2c, S2b-e). Cellular clustering was matched by increased chromatin accessibility over the corresponding cell-type specific TF binding sites such as illustrated for TBXT (Fig S2f), including over TEs found to be opened in these cellular subsets such as in axial mesodermal cells HERVH, which is also expressed in axial mesoderm of human gastrula (Fig 2d, S2g). Since TBXT expression is established at the primitive streak stage and maintained all along gastruloid formation^21^, it suggests that numerous HERVH loci are kept accessible by this TF throughout gastrulation. Intriguingly, despite the chromatin accessibility variation between cell types, we observed that a hundred TE subfamilies were maintained accessible at large (Fig S2h).

To investigate chromatin accessibility during later developmental stages we compared single-cell ATAC-seq data generated from 15 organs containing 54 different cell types derived from 12 to 17 weeks old fetuses^22^ with those obtained in pre-implantation embryo^4^ and *in vitro* derived and differentiated hESC^23^. First, we observed that chromatin was significatively more accessible than random in pre-implantation embryo, *in vitro* derived and differentiated hESC over TE subfamilies that remained accessible at fetal stages (Fig S2h). Second, these most significantly accessible TE subfamilies were largely primate-restricted (Fig 2e). Third, while LTR-derived sequences (ERVs) only contribute about 8% of the human genome through some 600’000 integrants, they accounted for one quarter of chromatin accessible loci in fetal tissues (Fig 2f). Fourth, profiles derived from single-cell chromatin accessibility of fetal organs revealed differential chromatin accessibility for distinct TE subfamilies, mirroring their germ layer-restricted expression (Fig 2g).

Together, these results suggest a model whereby chromatin at evolutionary recent TE loci, notably belonging to the ERV subclass, is rendered accessible in human embryo by the recruitment of pluripotency and gastrulation transcription factors, and then maintained opened during the subsequent steps of human development by lineage-specific transcription factors.

### Tissue-specific transcription factors control cell type-restriction of TE-derived enhancer activity

While many TEs expressed in pre-implantation embryo and embryonic stem cells are accessible and harbor binding sites for pluripotency factors, the patterns observed at later stages for EGA-induced TE and other subfamilies suggested regulation by cell type-specific transcription factors (TFs). To probe this hypothesis at large, we examined in a recently publish resource^24^ the transcriptional changes induced at TEs by individual overexpression of 328 transcription factors in epiblast-derived human embryonic stem cells (hESC) (Fig S3a-b). We found that whereas more than 200,000 TE different loci were deregulated in the sum of all these experiments, each TF significantly induced only a restricted set of TE subfamilies (Fig 3a, S3c-d). TEs previously noted to be transcribed during the minor and major waves of EGA, that is, HERVL and HERVK respectively, were activated by their known cognate activators DUX4 and KLF4 (Fig S3e), supporting the validity of our approach. Overexpression in hESCs of TFs considered as markers of particular germ layers such as GATA6 for the meso-endoderm and SOX17 for endoderm and PGCs induced the TE subfamilies noted to be expressed in the corresponding cells from gastrulating embryo (Fig 3b). Furthermore, while hESC overexpression of the meso-endoderm-specific GATA6 activated both LTR5Hs and LTR5B-derived HERVK, the endoderm/PGC-specific SOX17 only induced LTR5Hs-derived HERVK, mimicking the pattern observed in gastrulating embryos (Fig S3f).

**Figure 3.**
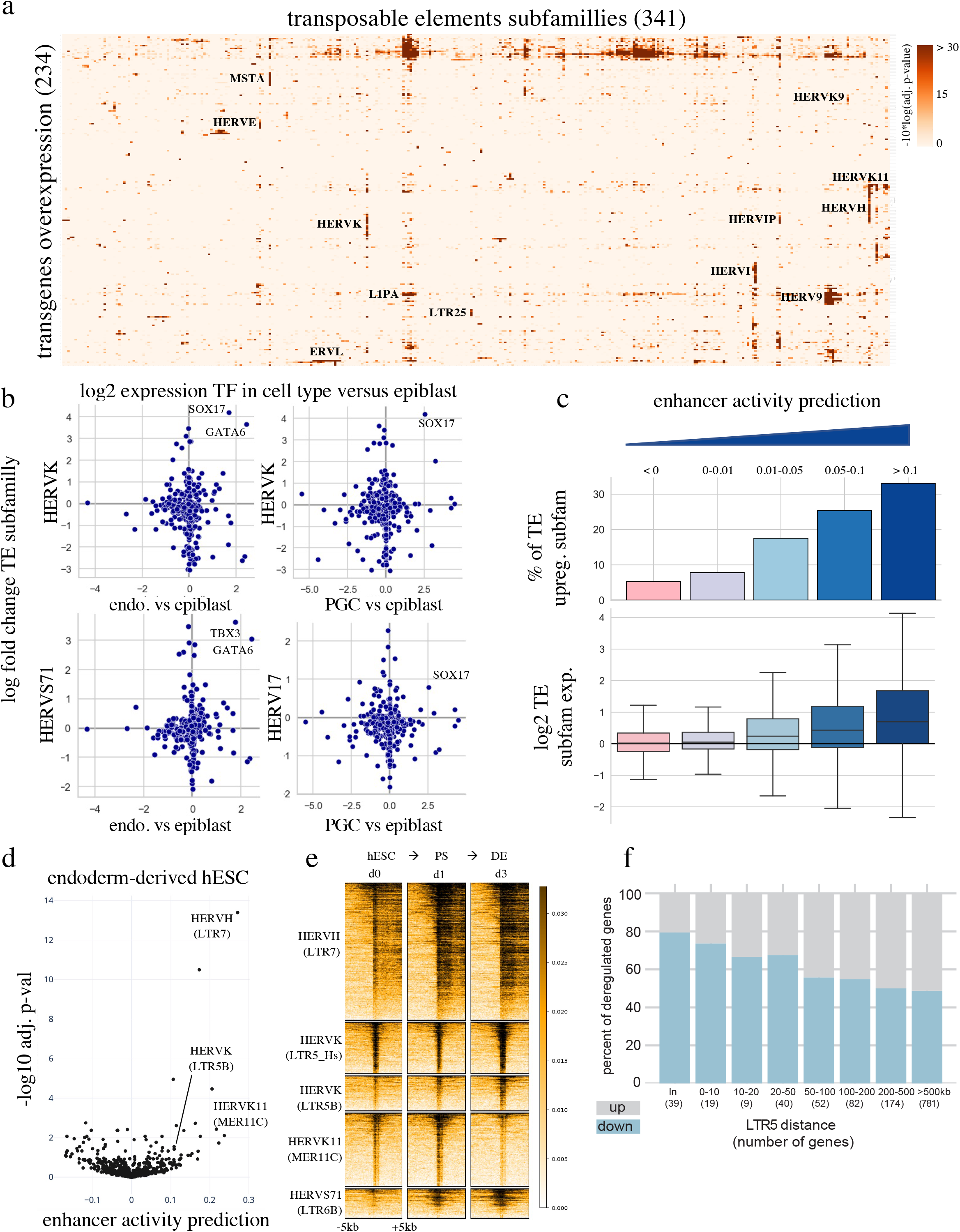
Evolutionary recent TEs act as cell-type-specific enhancers during human gastrulation and fetal development. **a**, Enrichment analysis of TE expression induced by transcription factor (TF) overexpression in hESC. Red color intensity corresponds to the enrichment (−10*log10[adjusted p-value]) of significantly up-regulated TEs over-representation for a given TE subfamily, re-analyzed from ^24^; only transgene overexpression conditions and TE subfamilies with an overrepresentation of increased expression (adjusted p-value < 0.05) among TEs expressed in each condition at least once are shown. **b**, Scatter plot illustrating the coupling between germ layer-specific TFs and TE subfamilies. y-axis, TE subfamily log2 fold change expression of intergenic subfamily (excluding all read overlapping a TEs in an exon, an intron and /-10kb of protein coding gene bodies) induced by overexpressed TF in hESC; x-axis, log2 fold change expression of these TFs in human gastrula versus epiblast cells **c**, Enhancer activity prediction of TE subfamily correlates with their transcriptional activation. Enhancer activity prediction was established for all TE subfamilies in all overexpressed transgenes conditions and grouped based on their activity values from the lowest to the highest (≤ 0, 0-0.01, 0.01-0.05, 0.05-0.1, >0.1); bar plots represent the % of transcriptionally induced TE subfamilies (adjusted p-value < 0.05, at the subfamily level of intergenic TEs) in each category; boxplots represent log2 fold TE subfamily add-up of normalized read counts expression change. **d**, TE-derived enhancer activity prediction upon endoderm differentiation. Representation of enhancer activity prediction for all TE subfamilies after comparing the transcriptome of hESC and hESC-derived endodermal cells after 3 days of differentiation; x-axis represent the activity value and the y-axis the -log10 adjusted p-value. **e**, H3K27ac enrichment over TE subfamily during hESC-derived endodermal differentiation. Black intensity correlates to H3K27ac ChIP-seq signal +/-10kbp around all TEs from a named subfamily. **f**, Impact of LTR5-targeting CRISPRi on gene expression during endodermal differentiation. Number of up- and downregulated genes (p-value < 0.05) at an indicated distance from closest CRISPRi-targeted TE is shown (in: TE within a gene).

In order to establish a link between TE activation by TFs and the possible regulation of neighboring genes by TE-originating *cis*-acting influences, we developed an algorithm of enhancer activity prediction. We used a linear regression model to seek a correlation between the presence of specific TE subfamily members in the vicinity of deregulated genes, which we expressed as an enhancer activity prediction score (Fig S3g). To validate this approach, we inhibited simultaneously SVA-and LTR5Hs-embedded transcriptional units by dCAS9-KRAB (CRISPRi)-mediated repression in naïve hESC. We then calculated the enhancer activity prediction scores of TE subfamilies, and confirmed that those most significantly affected in this setting corresponded to SVAs and LTR5Hs-derived HERVK targeted by CRISPRi (Fig S3h). Conversely, KLF4 overexpression in primed hESC resulted in increased enhancer activity prediction score for LTR5Hs-derived HERVK (Fig S3i), correlating our previous observation that this TF binds to these units and induces their transcription and acquisition of the H3K27ac active chromatin mark^9,24^. We then applied our algorithm to the analysis of the 328 hESC-TF overexpression dataset. We proceeded to rank TE subfamilies according to their enhancer activity prediction scores in each condition, and found a strong correlation with both the percentage of up-regulated integrants and the overall level of expression induction within this family for a given TF (Fig 3c). We also calculated the enhancer activity prediction score of individual TE subfamilies in hESC-derived endodermal cells and found it to be significant for HERVK11, LTR5B-derived HERVK as well as HERVHs (Fig 3d), with our enhancer prediction scores also correlating with increased H3K27ac loading, chromatin accessibility and expression of these TEs (Fig 3e and S3j). To validate functionally this observation, we targeted the LTR5 consensus sequence with CRISPRi in hESC (Fig S3k), which we then differentiated in endoderm. This resulted in the downregulation of hundreds of genes located near the corresponding TE integrants in the cellular products of this differentiation, confirming that these TE-embedded regulatory sequences acted as enhancers in this setting (Fig 3f). Of note, 21 genes encoding for KRAB-containing zing finger proteins (KZFPs) stood amongst the hundred nearby genes downregulated upon LTR5 repression (Fig S3l), a finding of interest since many members of this large family of DNA-binding proteins are responsible for silencing TEs through H3K9me3 deposition, histone deacetylation and DNA methylation^25^.

### Primate specific cis-and trans-regulators partner up to shape transcription during human gastrulation

As a consequence of their expansion by gene and segment duplications, many *KZFP* genes are grouped in clusters notably found on human chromosome 19. We observed that *KZFP* genes located in the same genomic cluster are often of similar evolutionary age, and are generally surrounded by insertions of contemporaneous TE subfamilies (Fig 4a, S4ab). Moreover, we noted while *KZFP* genes expression globally decreases during differentiation, primate-restricted clusters display coordinated upregulation in specific cells of human gastrula (Fig S4cd).

**Figure 4.**
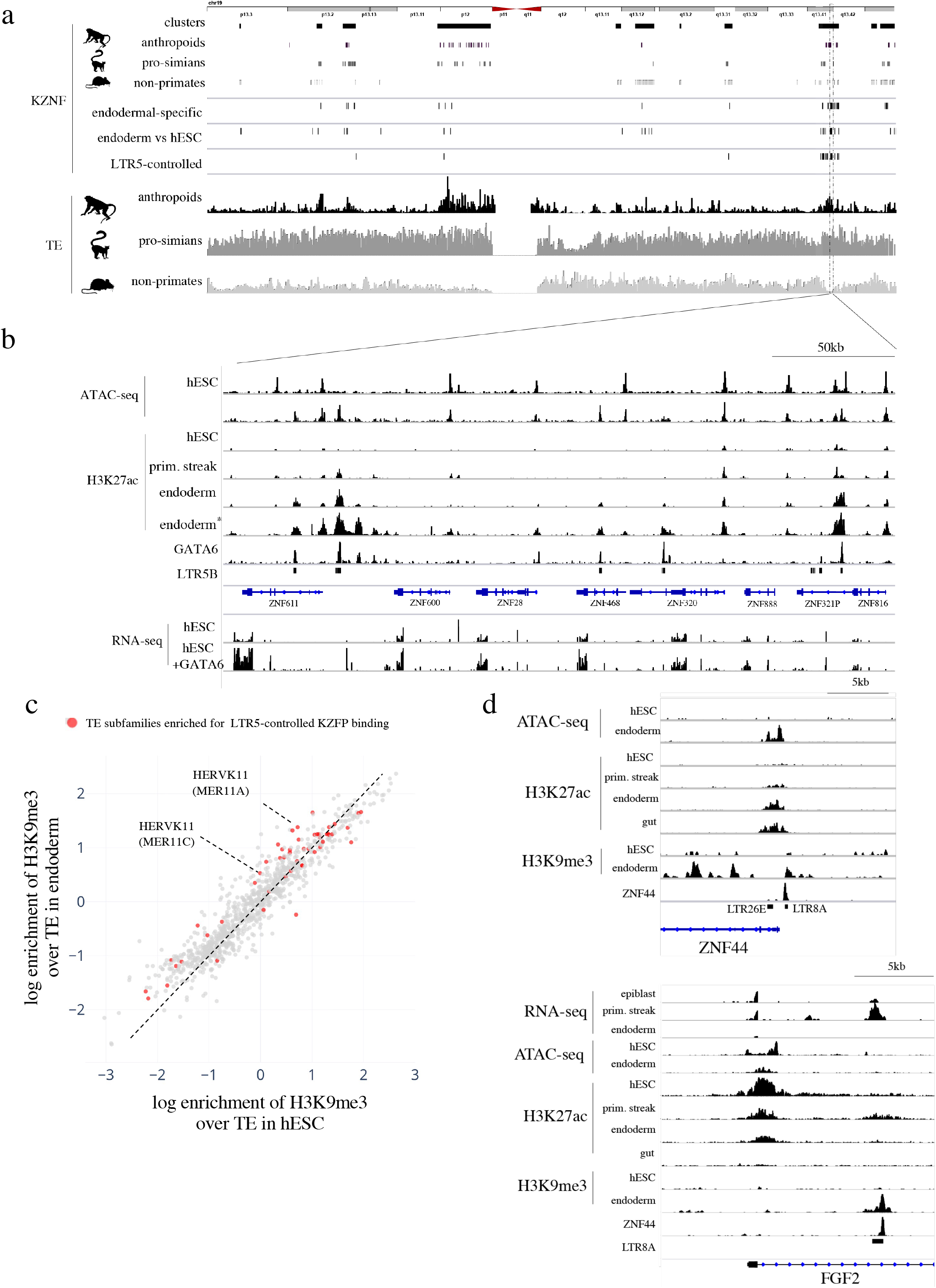
Primate specific cis- and trans-regulators partner up to control human gastrulation. **a**, Transcriptional KZFP regulation during endodermal differentiation, evolutionary age, and genomic distribution of TEs and KZFPs located on chromosome 19. Top, schematic representation of human chromosome 19 with KZFP gene clusters and individual KZFP. Middle, from top to bottom: KZFPs significantly upregulated in endodermal compared to other cells in human gastrula, in hESC-derived endoderm compared to hESC cells, and subjected to LTR5 control (as indicated by downregulation upon LTR5-targeting CRISPRi during endodermal differentiation with a p-value of 0.05). Bottom, TE density classified by evolutionary ages. **b**, Chromatin landscape of endodermal-specific KZFP cluster. Top, ATAC-seq and H3K27ac ChIP-seq profiles during endoderm differentiation at embryonic stem cell stage (hESC, day 0), primitive streak stage (prim. streak, day 1), and endodermal stage (endoderm, day 3) re-analyze from^53^; H3K27ac (endoderm*) and GATA6 ChIP-seq correspond to the same kinetic dataset than the ATAC-seq re-analyzed from^38^; bottom, RNA-seq performed in doxycycline-inducible GATA6 hESC line with (hESC+GATA6) or without (hESC) doxycycline during 48h, re-analyzed from ^24^. **c**, H3K9me3 differential enrichment in hESC *vs* hESC-derived endoderm. H3K9me3 TE subfamily enrichment was plotted for all subfamilies containing at least 10 integrants enriched in this mark in at least one condition; in red represent the TE subfamily significantly targeted by LTR5-controlled KZFPs. **d**, Dynamic control of FGF2 by LTR8A-derived enhancer and its KZFP controller ZNF44. Upper panel, genome browser visualization of *ZNF44* gene with from top to bottom ATAC-Seq, H3K27ac and H3K9me3 ChIP-seq profiles during endoderm differentiation from hESC (day 0), primitive streak stage (prim. streak, day 1), endodermal stage (endoderm, day 3) and gut stage (gut, day 7), re-analyzed from ^53^; ZNF44 ChIP-seq underneath is from^54^ and was performed in 293T. Lower panel, genome browser visualization of *FGF2* gene promoter, with from top to bottom single-cell human gastrula RNA-seq merged by cell-types of epiblast, primitive streak and endoderm, H3K27ac, and H3K9me3 ChIP-seq profiles during endoderm differentiation at embryonic stem cell stage (hESC, day 0), primitive streak stage (prim. streak, day 1), endodermal stage (endoderm, day 3), gut stage (gut, day 7), and ZNF44 ChIP-seq performed in 293T.

Most *KZFP* genes (19 genes, 17 of them primate-restricted) repressed upon LTR5 CRISPRi-mediated silencing during endodermal differentiation reside in one such cluster, which we found to be enriched in LTR5B inserts (Fig 4a). Eleven of these *KZFP* genes, which include *ZNF611, ZNF600, ZNF28, ZNF468* and *ZNF320* were more expressed in human gastrula endodermal cells and during *in vitro* endodermal differentiation of hESCs (Fig 4ab). Correspondingly, we determined that GATA6 bound directly to numerous LTR5B integrants during endodermal differentiation and induce LTR5B-derived HERVK and LTR5B-stimulated *KZFP* genes expression when overexpressed in hESC (Fig 4b). As well, the CRISPRa-mediated activation of LTR5 in NCCIT teratocarcinoma cells led to the induction of *KZFP* genes flanked by integrants belonging to this TE subset (Fig S4e).

Interestingly, we observed strong changes in the H3K9me3 landscape on TEs between hESC and hESC-derived endodermal cells and noted correlation with overlapping targets of LTR5-controlled KZFPs (Fig 4c). In particular, we observed LTR5-dependent increases in H3K9me3 enrichment over the transiently primitive-streak/nascent mesoderm expressed HERVK11 subfamily during *in vitro* endodermal differentiation (Fig 4c. S4fg).

This suggested a model whereby during gastrulation master TFs trigger a chain reaction by activating primate-restricted TEs and secondarily KZFPs of similar evolutionary age, which in turn repress these and other TE-based *cis*-acting regulatory elements, thus shaping the establishment of the various germ layers heterochromatin landscape. Lending credence to this model, we also found that repressing LTR5 integrants by CRISPRi triggered the upregulation of many mesoderm-specific genes in an experimental protocol of hESC-to-endoderm differentiation (Fig S4h). As more direct evidence of the impact of evolutionary intertwined TE-KZFPs on gastrulation, we first observed that during endodermal differentiation of hESCs, several LTR8A units become enriched for H3K9me3, whereas the LTR8A-targeting *ZNF44* displays increased accessibility and acetylation together with a nearby LTR26E-derived enhancer. Second, LTR8A-centered H3K9me3 accumulated in the vicinity of the *ZNF44* promoter itself, suggesting that this gene retro-controlled its expression (Fig 4d, upper panel). Finally, another H3K9me3-enriched LTR8 locus, near the *FGF2* promoter, serve as intronic enhancer in epiblast and primitive streak but is repressed in endoderm cells of human gastrula (Fig. 4d, lower panel). Of note, there is no such FGF2 transient expression in the mouse^19^, the genome of which contains neither LTR8 nor *ZNF44*.

Altogether, these data indicate that transcription networks at work during human early development, while driven by canonical transcription factors, are shaped by a partnership between primate-restricted TEs targeted by these activators and KZFPs countering their influences. As such, the regulome of human gastrulation and subsequent steps of fetal development displays a high level of species-specificity.

## DISCUSSION

Altogether, these data demonstrate that transcription networks at work during human early development, while orchestrated by canonical transcription factors, are shaped by a partnership between primate-restricted TE-embedded regulatory sequences (TEeRS) targeted by these activators and KZFP repressors countering their influences. As such, the regulome of human gastrulation displays a remarkable level of species-specificity. The broad expression of TEs such as HERVKs, HERVHs, SVAs and young LINE-1s during embryonic genome activation and in primordial germ cells^3,26^ is explained by their recognition by stem cell factors at play during these periods, such as SOX2, OCT4, NANOG or KLF4/17^7,9-13^. It fits with the biology of these TEs, new integrants of which have to seed the genome during the preimplantation period or germline formation to become inherited.

Our data indicate that TEeRS regulate subsequent steps of embryonic development in part through the recruitment of transcription factors active in germ layer determination, such as GATA6 and SOX17. More generally, it is well established that TEs can harbor binding sites for a wide array of transcription factors active in differentiated tissues, and the expression of some somatic genes, such as in the immune system, is driven by TEeRS^27-30^. Additionally, primate-specific TEeRS are major contributors to *cis*-regulatory innovations in human embryonic stem cells and adult liver^33,34^. The *cis*-regulatory influences of these recently emerged TEsRS on human fetal development suggest that, in spite of the evolutionary constraints proposed in the hourglasses developmental model^36^, all stages of this process are subjected to regulatory innovation.

Still, how the presence of binding sites for transcription factors typically active in differentiated tissues, which is predicted to promote neither the spread nor the inheritability of TEs, came to be selected through evolution is the object of much speculation^31,32^. However, we note here that primate-restricted ERVs are vastly overrepresented amongst TEs that harbor somatic TF binding sites, are expressed during gastrulation and display an open chromatin state during fetal development, consistent with the observation that many act as enhancers or promoters in developing or differentiated organs^35^. As ERVs are derived from exogenous retroviruses that once replicated in somatic tissues and were for many endowed with oncogenic properties favoring their expansion and persistence, it maybe that the diversity of TF binding sites harbored by ERVs is just a consequence of their ancestry. But how was this feature maintained in evolution? Our finding that TEs activated during either EGA or gastrulation stimulate the transcription of *KZFP* genes, the products of which in turn repress these TEs and confer germ layer-specificity to their transcriptional influences, strongly suggests that KZFPs, rather than just the host side of an evolutionary arms race^37^, were instrumental in allowing for the preservation and exploitation of the broad regulatory potential of TEs in higher vertebrates.

## Materials and Methods

### Embryonic Stem cell culture and differentiation

H9 human ESC line were maintained in mTSER plus on Matrigel and were passaged using TryplE in single cells. Endodermal differentiation was performed as in ^38^. Briefly, hESC were passaged at 80k cells/cm2 density in 12-well Matrigel-coated plate. When cells reached 80 – 90% confluence at 48-76h post-splitting, endodermal differentiation was initiated with 100 ng/ml of Activin A for 3 days, 5 μM GSK-3 inhibitor (CHIR-99021) to activate the WNT pathway for the first day, and 0.5 μM for the second day. Gastruloid differentiation was performed as in ^21^. Briefly, hESC were passaged at 40k cells/cm2 in mTSER density and at 48-76h post-splitting, mTSER was replace with Nutristem media with 3 μM CHIR-99021. After 24h cells were split in single-cell passaging with TryplE, embryoid bodies were formed with 800 cells per well of low adherence 96-well plate Cell Star in E6 media supplemented with 3 μM CHIR-99021 and 10 μM Rock inhibitor for 18 hours, then replace with E6 twice to be harvest 72h.

### CRISPRi experiments

sgRNA designed was perform taking Dfam consensus of LTR5B common sequence. Specificity was predicted with CRISPOR software (Haeussler et al., 2016). Human Embryonic Stem cells H9 media were transduced with dCAS9-KRAB lentiviral vector, selected and maintain in puromycin (0.25 μg/mL) for 5 to 10 days before endodermal differentiation experiments.

### ChIP-seq

Cells were cross-linked for 10 minutes at room temperature by the addition of one-tenth of the volume of 11% formaldehyde solution to the PBS followed by quenching with glycine. Cells were washed twice with PBS, then the supernatant was aspirated and the cell pellet was conserved in -80°C. Pellets were lysed, resuspended in 1mL of LB1 on ice for 10 min (50 mM HEPES-KOH pH 7.4, 140 mM NaCl, 1 mM EDTA, 0.5 mM EGTA, 10% Glycerol, 0.5% NP40, 0.25% Tx100, protease inhibitors), then after centrifugation resuspend in LB2 on ice for 10 min (10 mM Tris pH 8.0, 200 mM NaCl, 1 mM EDTA, 0.5 mM EGTA and protease inhibitors). After centrifugation, resuspend in LB3 (10 mM Tris pH 8.0, 200 mM NaCl, 1 mM EDTA, 0.5 mM EGTA, 0.1% NaDOC, 0.1% SDS and protease inhibitors) for histone marks and SDS shearing buffer (10 mM Tris pH8, EDTA 1mM, SDS 0.15% and protease inhibitors) for transcription factor and sonicated (Covaris settings: 5% duty, 200 cycle, 140 PIP, 20 min), yielding genomic DNA fragments with a bulk size of 100-300bp. Coating of the beads with the specific antibody and carried out during the day at 4°C, then chromatin was added overnight at 4°C for histone marks while antibody for transcription factor is incubated with chromatin first with 1% Triton and 150mM NaCl. Subsequently, washes were performed with 2x Low Salt Wash Buffer (10 mM Tris pH 8, 1 mM EDTA, 150mM NaCl, 0.15% SDS), 1x High Salt Wash Buffer (10 mM Tris pH 8, 1 mM EDTA, 500 mM NaCl, 0.15% SDS), 1x LiCl buffer (10 mM Tris pH 8, 1 mM EDTA, 0.5 mM EGTA, 250 mM LiCl, 1% NP40, 1% NaDOC) and 1 with TE buffer. Final DNA was purified with QIAGEN Elute Column. Up to 10 nanograms of ChIPed DNA or input DNA (Input) were prepared for sequencing. Library was quality checked by DNA high sensitivity chip (Agilent). Quality controlled samples were then quantified by picogreen (Qubit® 2.0 Fluorometer, Invitrogen).

Sequenced reads were aligned to the reference human genome hg19 with bowtie2 ^39^. MACS 1.4 ^40^ was used for peak calling. Peaks were merged using bedtools ^41^. FeatureCounts ^42^ was used to count uniquely mapped reads (MAPQ>10) on the peaks. Library size correction was performed using the TMM method as implemented in the limma package of R, using the total number of aligned reads as size factor.

Differential analysis on the uniquely mapped counts between conditions was performed with voom (REF). Heatmaps and profile averages were calculated using deeptools ^43^ over 5kb windows around the peak/repeat center from bigwigs. Enrichment analysis over TE subfamilies was performed with HOMER software^44^.

### RNA-seq

Total RNA from cell lines was isolated with NucleoSpin™ RNA Plus kit (Machery-Nagel). cDNA was prepared with Maxima Reverse Transcriptase (Thermo Scientific). Sequencing libraries were performed with Illumina Truseq Stranded mRNA LT kit. Reads were mapped to the human (hg19) genome using Hisat2 ^45^. Counts on genes and TEs were generated using featureCounts ^41^. To avoid read assignation ambiguity between genes and TEs, a gtf file containing both, genes and TEs was provided to featureCounts. For repetitive sequences, an in-house curated version of the Repbase database was used (fragmented LTR and internal segments belonging to a single integrant were merged). Only uniquely mapped reads were used for counting on genes and repetitive sequences integrants. TEs overlapping exons or that did not have at least one sample with 3 reads were discarded from the analysis. Normalization for sequencing depth has been done for both, genes and TEs, using the counts on genes as library size using the TMM method as implemented in the limma package of Bioconductor ^46^. Finally, for each transgene, differential gene expression analysis was performed using Voom ^47^ as it has been implemented in the limma package of Bioconductor and assessing only genes (or TEs) that had 3 reads in at least one sample. A gene (or TE) was considered to be differentially expressed when the fold change between groups was bigger than 2 and the p-value was smaller than 0.05. A moderated t-test (as implemented in the limma package of R) was used to test significance. P-values were corrected for multiple testing using the Benjamini-Hochberg’s method. For counting on TE subfamilies, we added up the the reads on the repetitive sequences without filtering out for multi-mapped reads and added them up per subfamily.

### Enhancer activity estimation

To identify TE subfamilies exerting putative enhancer activities directly from RNA-seq data, we modeled treatment vs control deviations in gene expression of protein coding genes as a linear combination of occurrences of nearby (within 50kb of the TSS) TE subfamily integrants. We excluded TEs overlapping exons and TE subfamilies that colocalized less than 150 times near the promoters of protein coding genes. The coefficients of this linear regression problem represent deviations in putative enhancer activities of TE subfamilies. Similar models were proposed to infer the activity of DNA motifs from gene expression data^48^. To find TE subfamilies with significant cis-regulatory activities, we performed null significance hypothesis testing on the linear regression coefficients and accounted for multiple testing using the Benjamini Hochberg procedure.

### ATAC-seq analysis

Reads were aligned with bowtie2. MT reads were removed before peak calling. Peak calling was done with MACS2 with FDR cutoff of 0.01 and q-value < 10^e-5^, and using the --bampe option when PE reads.

### Single-cell multi-omic

Cellranger-arc^49^ was used to obtain counts on genes and peaks using default parameters. The hg38 reference genome provided by cellranger-arc was used.

### Single-cell RNA-seq analysis

For human embryo dataset, counts were obtained using cellranger^50^ using a GTF of hg19 that contained both, genes and TEs. Only uniquely mapped reads on genes that were expressed in at least 1% of the samples and in minimum 3 cells for TEs were kept. Then, for TEs not overlapping exons, counts were added up at subfamily level. Cells with less than 200 features and more than 25% of mitochondrial reads were removed. Seurat’s SCTransform ^51^ was used to normalize the data and correct for mitochondrial percentage and total number of reads biases. For embryonic-body time course differentiation, idem except that only cells with more than 20% of mitochondrial reads were removed.

### Single-cell ATAC-seq analysis

Single-cell ATAQ-seq peaks data were downloaded from the Atlas Of Chromatin Accessibility During Development(https://atlas.brotmanbaty.org/bbi/human-chromatin-during-development/,^22^. TE family enrichment significance for chromatin accessibility were assess by random permutations: First, for each TE sub family, the total number of detected peaks for a given cell type on the selected TE sub family was computed using R GenomicRanges library. Then, TE sub families were randomly shuffled 10 times using bedtools with options –chrom and – noOverlapping and the total number of peaks for each permutation was computed. The fold enrichment of the significant subfamilies was then plotted on a heatmap using R heatmap2 function for each cell type.

### TE density profile

TE bigwig densities were computed using python. First, TEs, depending of their subfamily evolutionary ages, were extracted in bed format from the TE database and converted to bedgraph using the genomecov routine of BedTools. Then, bedgraph signals were smoothed using a rectangular window of 10kb and written in bigwig format using the pyBigWig python library.

## Data availability

Single-cell multi-omics of gastruloid, RNA-seq of endodermal differentiated hESC with or without LTR5-targeting sgRNA generated in this study were deposited on Gene Expression Omnibus (GEO) database under accession number GSE181120. All other genomic datas of this study: from GEO database: GSE140021 for the 10X single-cell RNA-seq of hESC-derived embryonic body time course; ATAC-seq of purified PGC during hESC-derived embryonic body differentiation: GSE120648. ChIP-seq and ATAC-seq from endodermal differentiation GSE117136 and GSE52657. ATAC-seq of naïve and primed hESC from GSE130418. Single-cell ATAC-seq from fetal organ: https://descartes.brotmanbaty.org/. GSE124128 for the H3K9me3 profile upon CRISPRa of LTR5 in NCCIT. E-MTAB-9388 for the SMART-seq single-cell RNA-seq of human gastrula from EBI database. DRA006296 for the overexpression dataset of TF in hESC from DDBJ database.

## Author information

### Author notes

These authors contributed equally: Julien Pontis and Didier Trono

### Contributions

J.P. and D.T. conceived the study and wrote the manuscript; J.P. designed and performed all experiments with the technical help of S.O.; J.P., C.P, E.P., D.G and J.D. completed the bioinformatics analyses; A.M. and M.L. provided valuable advice for gastruloid experiments.

## Ethics declarations

Human ESC usage has been approved by the Swiss Federal Office of Public Health, the Canton of Vaud Ethics Committee (Autorization Number R-FP-S-2-0009-0000) and registered in the European Human Pluripotent Stem Cell Registry (hPSCreg).

## Acknowledgments

We greatly thank Shankar Srinivas and Antonio Scialdone for providing single-cell expression raw data of the human gastrula. We also thank the EPFL Genomics core facilities and the University of Lausanne Genomic Technologies Facility for help with sequencing. We finally thank Alexandre Mayran for critical reading of the manuscript. This study was supported by grants from the Swiss National Science Foundation and the European Research Council (KRABnKAP, no. 268721; Transpos-X, no. 694658) to D.T.; by fellowships from the EPFL/Marie Skłodowska-Curie Fund, the Association pour la Recherche sur le Cancer (ARC), and the Fondation Bettencourt to J.P.

## Figure legends

**Figure S1 for Fig 1. Cell-type specific expression of primate-restricted TEs during human gastrulation**.

**a**, Cellular composition of human gastrula. UMAP based on single-cell gene expression from human gastrula. Colors represent a more detail cell subtypes identify in^19^. **b**, Age distribution of expressed intergenic TEs in human gastrula. Each TE subfamily (excluding DNA transposons) was restricted to a specific evolutionary age category, and the percentage of expressed TE integrants in each was plotted. We excluded any TEs overlapping coding gene body and up/down-stream TE, thus removing TE expressed due to readthrough or gene transcript inclusion. **c**, Cell-type-specific expression of TE subfamilies; boxplots with each dot representing the TE subfamily normalized expression in one cell. Cells were grouped in boxplot corresponding to one cell type of human gastrula sub-clustering: epiblast, primitive streak (prim. streak), primordial germ cells (PGC), nascent mesoderm (na.), emergent mesoderm (em.), advance mesoderm (adv.), axial mesoderm (axial), yolk mesoderm (yolk), definitive endoderm (def. endo.), definitive endoderm non-proliferative (def. endo*), hypoblast (hypo.), yolk endoderm (yolk), ectoderm (ectoderm), hemogenic endothelium (hemo-endo.), myeloid progenitor (mye-prog.), erythro-myeloid progenitor (ery-my p.), blood progenitor (blood prog.), erythrocytes (erythro.). **d**, Cell type-specific expression of TE subfamilies; heatmap of the -log10 adjusted p-value of cell type specificity of TE subfamily expression; only TE subfamilies with adjusted p-value < 10^e-5^ were plotted.

**Figure S2 for Fig 2. TEs are controlled by tissue-specific transcription factors**

**a**, Cell type-specific expression of transcription factors during embryoid body differentiation. Each plot represents a UMAP defined on single-cell gene expression during differentiation of hESC into embryoid bodies over 5 days re-analyzed from^20^; hESC corresponds to day 0; primitive streak (prim. streak) to days 1-2 with TBXT expression; cells at days 2-5 are stratified in PGC expressing NANOS3, NANOG, and SOX17, endoderm expressing SOX17 and GATA6, and mesoderm expressing GATA6 only. Color scale corresponds to level of relative TE subfamily expression based on normalized read counts. **b**, Gastruloid formation and nuclei isolation. Top, picture of pooled elongated gastruloids used for the multi-omic experiment; bottom, trypan blue of purified nuclei used for the multi-omic experiment. **c**, UMI (unique molecular identifier) distribution of gene expression and transposase cut site counts of chromatin accessibility in each cluster defined by expression in single-nuclei RNA-seq and ATAC-seq. Clusters 2 and 8 contained lower amount of RNA and/or ATAC-seq UMI/cut sites, hence were ignored for the rest of the analysis; clusters 1 and 4 look similar to cluster 6 but additionally express different cycling genes. **d**, UMAP of single-cell clustering of gastruloids based on gene expression; each color represents a different cluster illustrated in S2c. **e** Violin plot of expression level of indicated cell type-specific transcription factors each cluster from S2c; cell-type specific transcription factors were selected based on cluster-specificity of expression and DNA binding motif enrichment for their chromatin accessibility. **f**, Relative expression (y-axis) and motif enrichment at accessible chromatin (x-axis) of cell type-specific transcription factors in axial mesodermal cells (cluster 3) comparted with other clusters. **g**, Genome browser of TBXT binding (top) and merged single-cell chromatin accessibility at an HERVH integrant in hESC-derived mesoderm (from ^52^) (cluster 3) with early paraxial cells as a negative control (cluster 6). **h**, Heat map of relative chromatin accessibility over indicated TE subfamilies compared to random genomic distribution, using peaks obtained from DNAse-seq in morula, ATAC-seq in primed and naïve hESC, primitive-streak, endoderm and PGC derived hESC, as well as from merged single-cell ATAC-seq of hESC-derived gastruloids and fetal organs, re-analyzed from ^4,22,23^.; 171 TE subfamilies are represented, based on having in at least one condition 50 accessible integrants with significant overlap (p-value < 0.05), with 2 fold change enrichment over random genome coverage.

**Figure S3 for Fig 3. Evolutionary recent TEs act as cell-type-specific enhancers during human gastrulation and fetal development**.

**a**, Design of TFs overexpression in hESC experiment performed in ^24^. **b**, Overexpressed TF subtypes. Each bar represents the number of proteins harboring indicated domain, with fraction in blue indicating those overexpressed in ^2 4^. **c**, Sum of total, expressed and differentially expressed genes and TEs upon overexpression of 234 TFs in hESC (with adjusted p-value <0.05). **d**, Number of TE subfamilies deregulated per overexpressed TF; each bar plot represents the number of TFs inducing number TE subfamilies indicated on x-axis (adjusted p-value < 0.05 for significantly up-regulated TEs over-representation in a TE subfamily). **e**, Scatter plot illustrating coupling between EGA-induced TFs and TEs. y-axis, log2 fold TE subfamily add-up of normalized read counts expression change; x-axis, number of up-regulated TE integrants from that subfamily (2-fold with adjusted p-value <0.05). **f**, Scatter plot illustrating correlation between germ layer-specific TEs and TFs. y-axis, log2 fold TE subfamily add-up of normalized read counts expression induced by overexpressed transcription factors in hESC; x-axis, log2 fold change expression of these transcription factors in human gastrula versus epiblast cells. **g**, TE subfamily-derived enhancer activity estimation schema. Regressive correlation is applied based on each coding gene expression level (Ex) and neighboring representation (100kb around TSS) of each TE subfamilies (T) in each TF-overexpressing hESC (E), allowing estimation of each TE subfamily relative enhancer activity (A). **h**, T-statistical tests of TE subfamilies activity around deregulated genes upon CRISPRi-targeting SVA/LTR5Hs in naïve hESCs. **i**, Activity of LTR5Hs subfamily in the presence or absence of KLF4 overexpression in primed hESCs from^9^ in the left panel and^24^ in the right panel at 24h or 48h overexpression. **j**, Log2 number of expressed TE integrants induced upon hESC-derived endodermal differentiation; p-value enrichment is represented for each presented subfamily. **m**, Predicted number of targeted LTR5 by CRISPRi. **k**, Gene Ontology of nearby LTR5-controlled genes. All down-regulated genes upon LTR5-targeting CRISPRi in a 100kbp window from LTR5 were selected; 2D-pie represents the proportion of 21 KZFN genes among the 107 down-regulated genes.

**Figure S4 for Fig 4. Primate specific cis- and trans-regulators partner up to control human gastrulation**

**a**, Evolutionary young *KZFP* gene clusters are enriched in contemporary TEs. Top, boxplot of *KZFP* genes evolutionary ages within 11 clusters (C1 to C11) on chromosome 19 or elsewhere in the genome (others); bottom, boxplot of evolutionary ages of TE subfamilies enriched in these *KZFP* gene clusters (p-value < 10^e-4^). **b**, Heatmap of TE subfamily enrichment in *KZFP* gene clusters. Yellow intensity is proportional to significance (−10*log10(p-value)); only TE subfamily with a p-value enrichment of 10e-4, a fold change of 2 and at least 5 integrant inside one cluster were represented. **c**, Heatmap of *KZFP* genes expression in human gastrula re-analyzed from^19^. Yellow intensity is proportional to log normalized counts; only *KZFP* genes with at least 2 normalized counts were plotted. **d**, *KZFP* genes expression in human gastrula. Each boxplot represents log2 gene expression fold change of one cell type over the others. Top panel shows the expression of all *KZFP* genes; middle and bottom panels depict *KZFP* genes in clusters 4 and 9, respectively. **e**, LTR5-controlled *KZFP* genes vs other genes during endodermal differentiation are activated upon CRISPRa in NCCIT cell line. Left, boxplots of *KZFP* genes significantly (down) or not (others) down-regulated upon LTR5-mediated repression during endodermal differentiation; on the right are presented the fold change upon LTR5-mediated activation in NCCIT cell line of the *down* and *others* KZFPs re-analyzed from^55^. **f**, H3K9me3 loss at HERVK11 upon LTR5-mediated repression during endodermal differentiation. Panel represent H3K9me3 ChIP-qPCR upon LTR5 repression at this same HERVK11 elements; t.test were performed with non-significant (n.s) or < 0.005 (***) results on biological quadruplicates. **g**, H3K9me3 profile at HERVK11 elements during endodermal differentiation. Panel represent H3K9me3 ChIP-seq profile in hESC and endoderm re-analyzed from Encode datasets. **h**, Cell type-specific genes of human gastrula up-regulated upon LTR5 repression during endodermal differentiation. Genes significantly up-regulated (p-value < 0.05) upon LTR5-targeted CRISPRi during endodermal differentiation were selected and crossed with cell type-specific genes from human gastrula (p-value < 0.05); y-axis represent log2 fold change expression for all human gastrula cell type-specific genes versus epiblast cells.

## References

1. Sundaram, V. & Wysocka, J. Transposable elements as a potent source of diverse cis-regulatory sequences in mammalian genomes. Philos Trans R Soc Lond B Biol Sci 375, 20190347 (2020).

2. Chuong, E. B., Elde, N. C. & Feschotte, C. Regulatory activities of transposable elements: from conflicts to benefits. Nat. Rev. Genet. 18, 71–86 (2017).

3. Göke, J. et al. Dynamic transcription of distinct classes of endogenous retroviral elements marks specific populations of early human embryonic cells. Cell Stem Cell 16, 135–141 (2015).

4. Gao, L. et al. Chromatin Accessibility Landscape in Human Early Embryos and Its Association with Evolution. Cell 173, 248–259.e15 (2018).

5. Liu, L. et al. An integrated chromatin accessibility and transcriptome landscape of human pre-implantation embryos. Nature Communications 10, 364–11 (2019).

6. Wu, J. et al. Chromatin analysis in human early development reveals epigenetic transition during ZGA. Nature Publishing Group 557, 256–260 (2018).

7. Kunarso, G. et al. Transposable elements have rewired the core regulatory network of human embryonic stem cells. Nat. Genet. 42, 631–634 (2010).

8. Fort, A. et al. Deep transcriptome profiling of mammalian stem cells supports a regulatory role for retrotransposons in pluripotency maintenance. Nat. Genet. 46, 558–566 (2014).

9. Pontis, J. et al. Hominoid-Specific Transposable Elements and KZFPs Facilitate Human Embryonic Genome Activation and Control Transcription in Naive Human ESCs. Cell Stem Cell 24, 724–735.e5 (2019).

10. Wang, J. et al. Primate-specific endogenous retrovirus-driven transcription defines naive-like stem cells. Nature Publishing Group 516, 405–409 (2014).

11. Ohnuki, M. et al. Dynamic regulation of human endogenous retroviruses mediates factor-induced reprogramming and differentiation potential. Proceedings of the National Academy of Sciences of the United States of America 111, 12426–12431 (2014).

12. Grow, E. J. et al. Intrinsic retroviral reactivation in human preimplantation embryos and pluripotent cells. Nature 522, 221–225 (2015).

13. Barakat, T. S. et al. Functional Dissection of the Enhancer Repertoire in Human Embryonic Stem Cells. Stem Cell 23, 276–288.e8 (2018).

14. Haring, N. L. et al. ZNF91 deletion in human embryonic stem cells leads to ectopic activation of SVA retrotransposons and up-regulation of KRAB zinc finger gene clusters. Genome Res. 31, 551–563 (2021).

15. Greenberg, M. V. C. & Bourc’his, D. The diverse roles of DNA methylation in mammalian development and disease. Nature Reviews Molecular Cell Biology 20, 590–607 (2019).

16. Friedli, M. & Trono, D. The developmental control of transposable elements and the evolution of higher species. Annu. Rev. Cell Dev. Biol. 31, 429–451 (2015).

17. Guo, H. et al. The DNA methylation landscape of human early embryos. Nature Publishing Group 511, 606–610 (2014).

18. Smith, Z. D. et al. DNA methylation dynamics of the human preimplantation embryo. Nature Publishing Group 511, 611–615 (2014).

19. Tyser, R. C. V. et al. A spatially resolved single cell atlas of human gastrulation. bioRxiv 2020.07.21.213512 (2020). doi:10.1101/2020.07.21.213512

20. Chen, D. et al. Human Primordial Germ Cells Are Specified from Lineage-Primed Progenitors. Cell Rep 29, 4568–4582.e5 (2019).

21. Moris, N. et al. An in vitro model of early anteroposterior organization during human development. Nature Publishing Group 582, 410–415 (2020).

22. Domcke, S. et al. A human cell atlas of fetal chromatin accessibility. Science 370, (2020).

23. Pastor, W. A. et al. TFAP2C regulates transcription in human naive pluripotency by opening enhancers. Nature Cell Biology 20, 553–564 (2018).

24. Nakatake, Y. et al. Generation and Profiling of 2,135 Human ESC Lines for the Systematic Analyses of Cell States Perturbed by Inducing Single Transcription Factors. Cell Rep 31, 107655 (2020).

25. Ecco, G., Imbeault, M. & Trono, D. KRAB zinc finger proteins. Development 144, 2719–2729 (2017).

26. Tang, W. W. C. et al. A Unique Gene Regulatory Network Resets the Human Germline Epigenome for Development. Cell 161, 1453–1467 (2015).

27. Bourque, G. et al. Evolution of the mammalian transcription factor binding repertoire via transposable elements. Genome Res. 18, 1752–1762 (2008).

28. Sundaram, V. et al. Widespread contribution of transposable elements to the innovation of gene regulatory networks. Genome Res. 24, 1963–1976 (2014).

29. Ito, J. et al. Systematic identification and characterization of regulatory elements derived from human endogenous retroviruses. PLoS Genet. 13, e1006883 (2017).

30. Chuong, E. B., Elde, N. C. & Feschotte, C. Regulatory evolution of innate immunity through co-option of endogenous retroviruses. Science 351, 1083–1087 (2016).

31. Sundaram, V. & Wang, T. Transposable Element Mediated Innovation in Gene Regulatory Landscapes of Cells: Re-Visiting the ‘Gene-Battery’ Model. BioEssays 40, (2018).

32. Britten, R. J. & Davidson, E. H. Repetitive and Non-Repetitive DNA Sequences and a Speculation on the Origins of Evolutionary Novelty. Q Rev Biol 46, 111–138 (1971).

33. Trizzino, M. et al. Transposable elements are the primary source of novelty in primate gene regulation. Genome Res. 27, 1623–1633 (2017).

34. Jacques, P.-É., Jeyakani, J. & Bourque, G. The Majority of Primate-Specific Regulatory Sequences Are Derived from Transposable Elements. PLoS Genet. 9, e1003504 (2013).

35. Pehrsson, E. C., Choudhary, M. N. K., Sundaram, V. & Wang, T. The epigenomic landscape of transposable elements across normal human development and anatomy. Nature Communications 10, 5640 (2019).

36. Development, D. D. 1994. Temporal colinearity and the phylotypic progression: a basis for the stability of a vertebrate Bauplan and the evolution of morphologies through heterochrony. journals.biologists.com

37. Jacobs, F. M. J. et al. An evolutionary arms race between KRAB zinc-finger genes ZNF91/93 and SVA/L1 retrotransposons. Nature 516, 242–245 (2014).

38. Lee, K. et al. FOXA2 Is Required for Enhancer Priming during Pancreatic Differentiation. Cell Rep 28, 382–393.e7 (2019).

39. Langmead, B. & Salzberg, S. L. Fast gapped-read alignment with Bowtie 2. Nat. Methods 9, 357–359 (2012).

40. Zhang, Y. et al. Model-based analysis of ChIP-Seq (MACS). Genome Biol. 9, R137 (2008).

41. Quinlan, A. R. & Hall, I. M. BEDTools: a flexible suite of utilities for comparing genomic features. Bioinformatics 26, 841–842 (2010).

42. Liao, Y., Smyth, G. K. & Shi, W. featureCounts: an efficient general purpose program for assigning sequence reads to genomic features. Bioinformatics 30, 923–930 (2014).

43. Ramírez, F., Dündar, F., Diehl, S., Grüning, B. A. & Manke, T. deepTools: a flexible platform for exploring deep-sequencing data. Nucleic Acids Res. 42, W187–91 (2014).

44. Heinz, S. et al. Simple Combinations of Lineage-Determining Transcription Factors Prime cis-Regulatory Elements Required for Macrophage and B Cell Identities. Molecular Cell 38, 576–589 (2010).

45. Kim, D., Langmead, B. & Salzberg, S. L. HISAT: a fast spliced aligner with low memory requirements. Nat. Methods 12, 357–360 (2015).

46. Gentleman, R. C. et al. Bioconductor: open software development for computational biology and bioinformatics. Genome Biol. 5, R80 (2004).

47. Law, C. W., Chen, Y., Shi, W. & Smyth, G. K. voom: Precision weights unlock linear model analysis tools for RNA-seq read counts. Genome Biol. 15, R29 (2014).

48. Balwierz, P. J. et al. ISMARA: automated modeling of genomic signals as a democracy of regulatory motifs. Genome Res. 24, 869–884 (2014).

49. Satpathy, A. T. et al. Massively parallel single-cell chromatin landscapes of human immune cell development and intratumoral T cell exhaustion. Nature Biotechnology 37, 925–936 (2019).

50. Molè, M. A. et al. A single cell characterisation of human embryogenesis identifies pluripotency transitions and putative anterior hypoblast centre. Nature Communications 12, 3679 (2021).

51. Stuart, T. et al. Comprehensive Integration of Single-Cell Data. Cell 177, 1888–1902.e21 (2019).

52. Faial, T. et al. Brachyury and SMAD signalling collaboratively orchestrate distinct mesoderm and endoderm gene regulatory networks in differentiating human embryonic stem cells. Development 142, 2121–2135 (2015).

53. Loh, K. M. et al. Efficient endoderm induction from human pluripotent stem cells by logically directing signals controlling lineage bifurcations. Cell Stem Cell 14, 237–252 (2014).

54. Imbeault, M., Helleboid, P.-Y. & Trono, D. KRAB zinc-finger proteins contribute to the evolution of gene regulatory networks. Nature Publishing Group 543, 550–554 (2017).

55. Fuentes, D. R., Swigut, T. & Wysocka, J. Systematic perturbation of retroviral LTRs reveals widespread long-range effects on human gene regulation. Elife 7, (2018).

