## Supplementary figures for "Primate-specific *cis*- and *trans*-regulators shape transcriptional networks during human development"

**Fig. S1: Cell-type Specific Expression of Primates TEs during Human Gastrulation.**

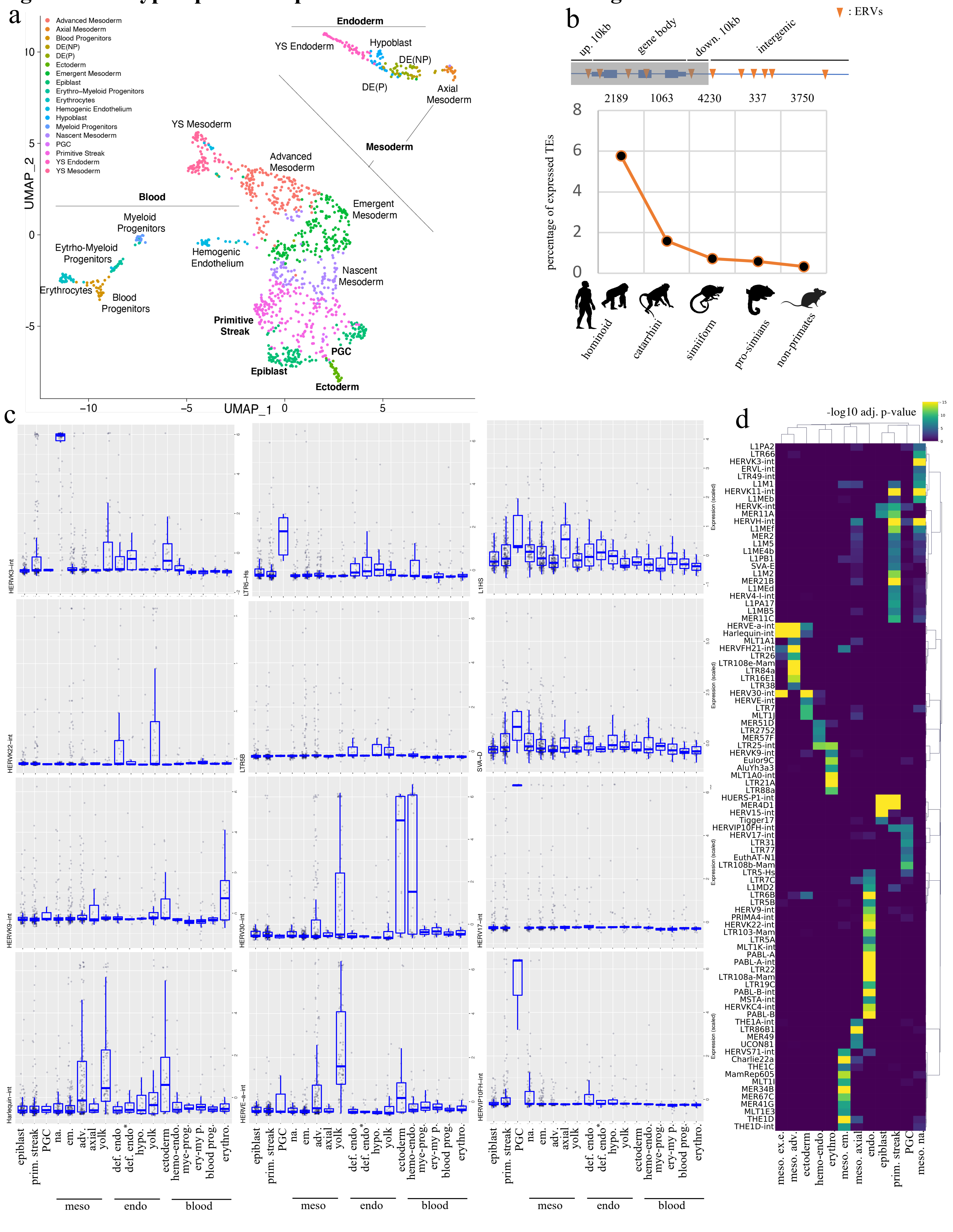

### Fig. S2: Evolutionary recent TEs maintain their cis-regulatory potential during human gastrulation and fetal development

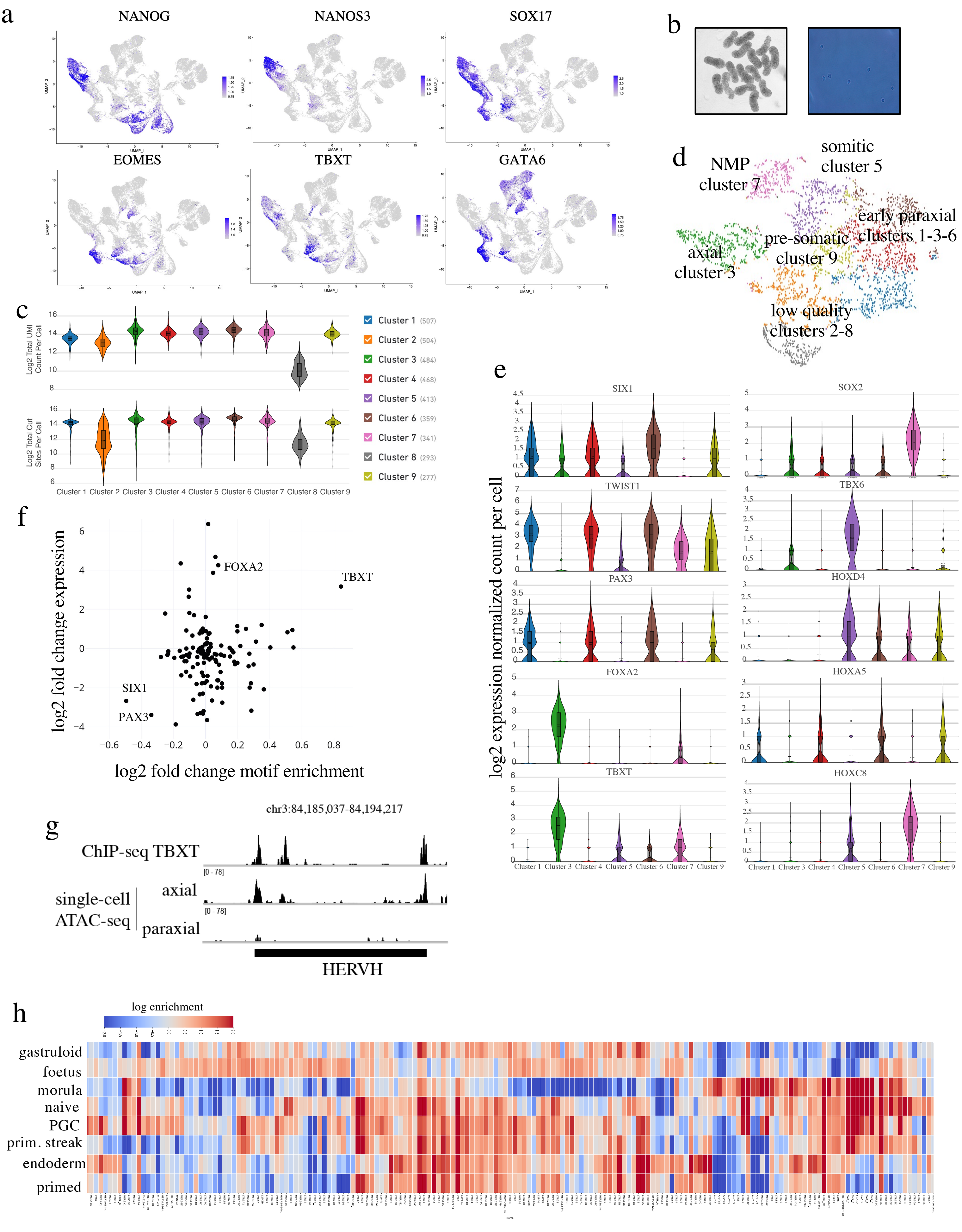

### Fig. S3: Tissue-specific transcription factors control cell-type-specificity of TE-derived enhancer activity

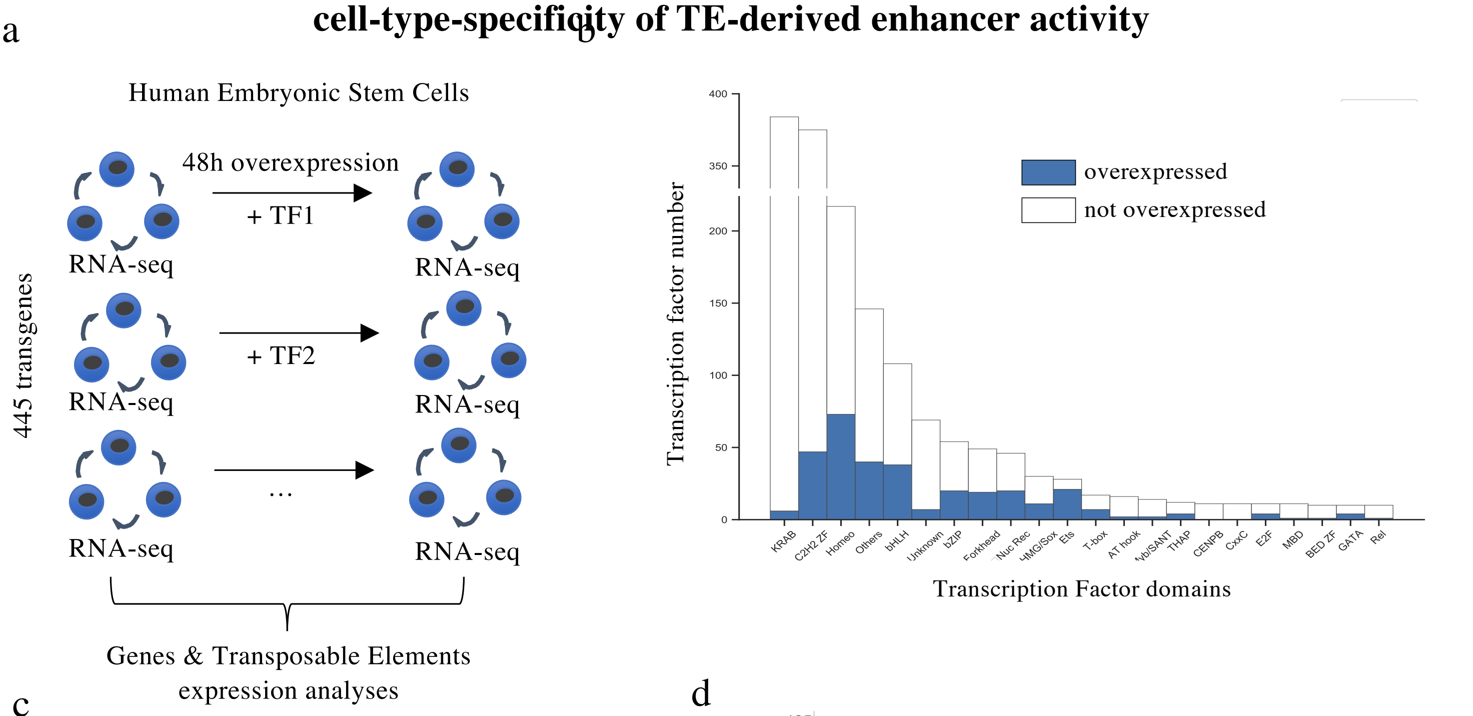


c

| | coding genes | transposable elements |
|---|---|---|
| total | 20'327 | 4'570'939 |
| expressed | 18'817 | 397'163 |
| deregulated | 18'428 | 216'160 |

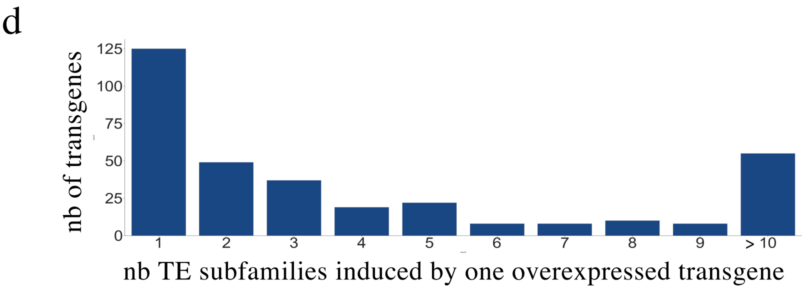


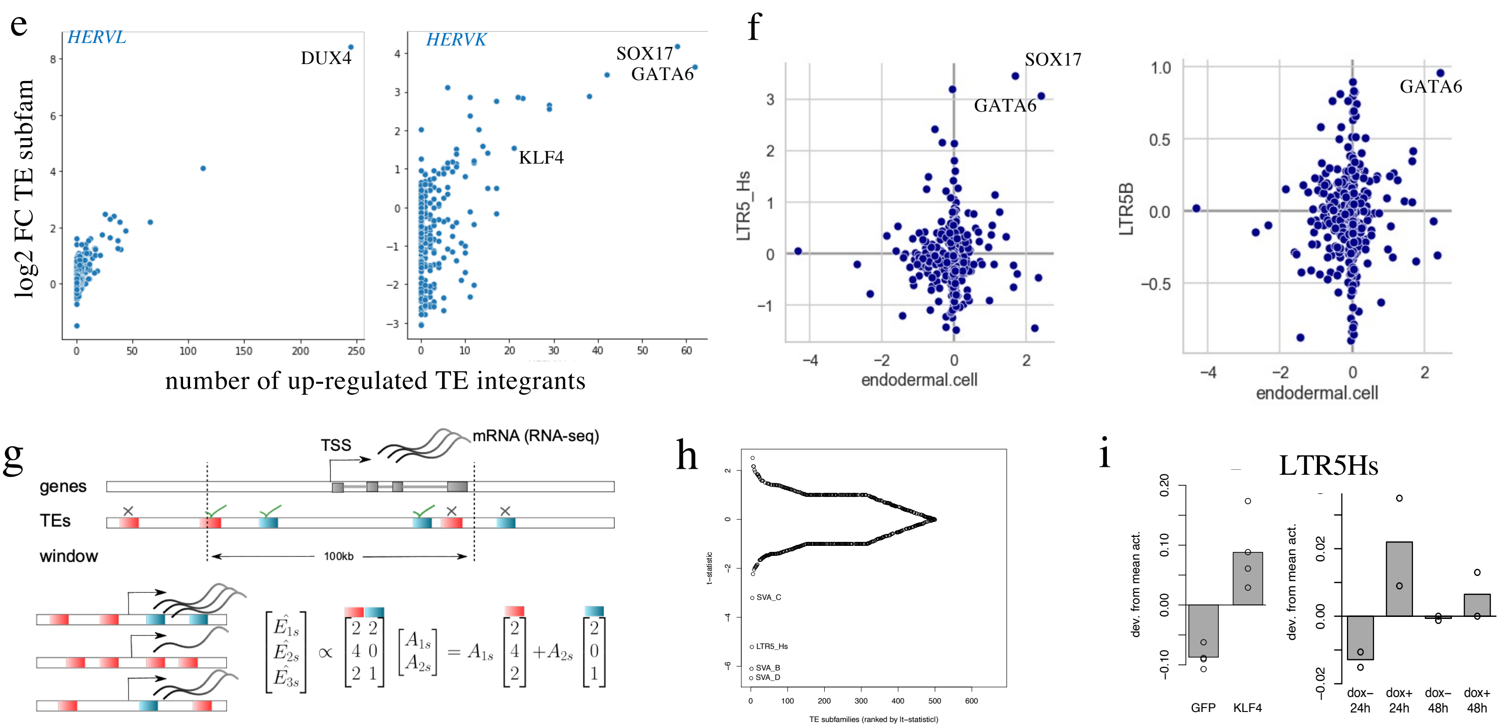


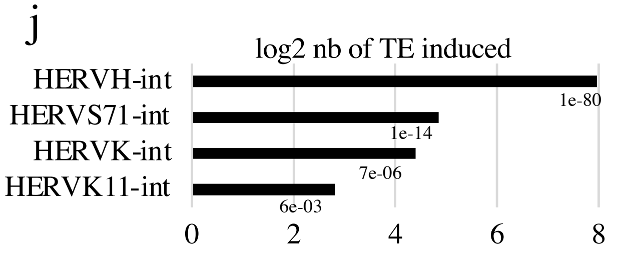


k

| TE subfamilly name | number of integrants | predicted targets |
|---|---|---|
| LTR5_Hs | 645 | 483 |
| LTR5B | 431 | 178 |
| LTR5A | 265 | 1 |
| LTR5 | 20 | 2 |

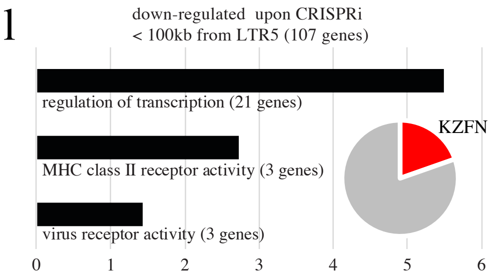

### Fig. S4: Primate specific cis- and trans-regulators partner up to control human gastrulation

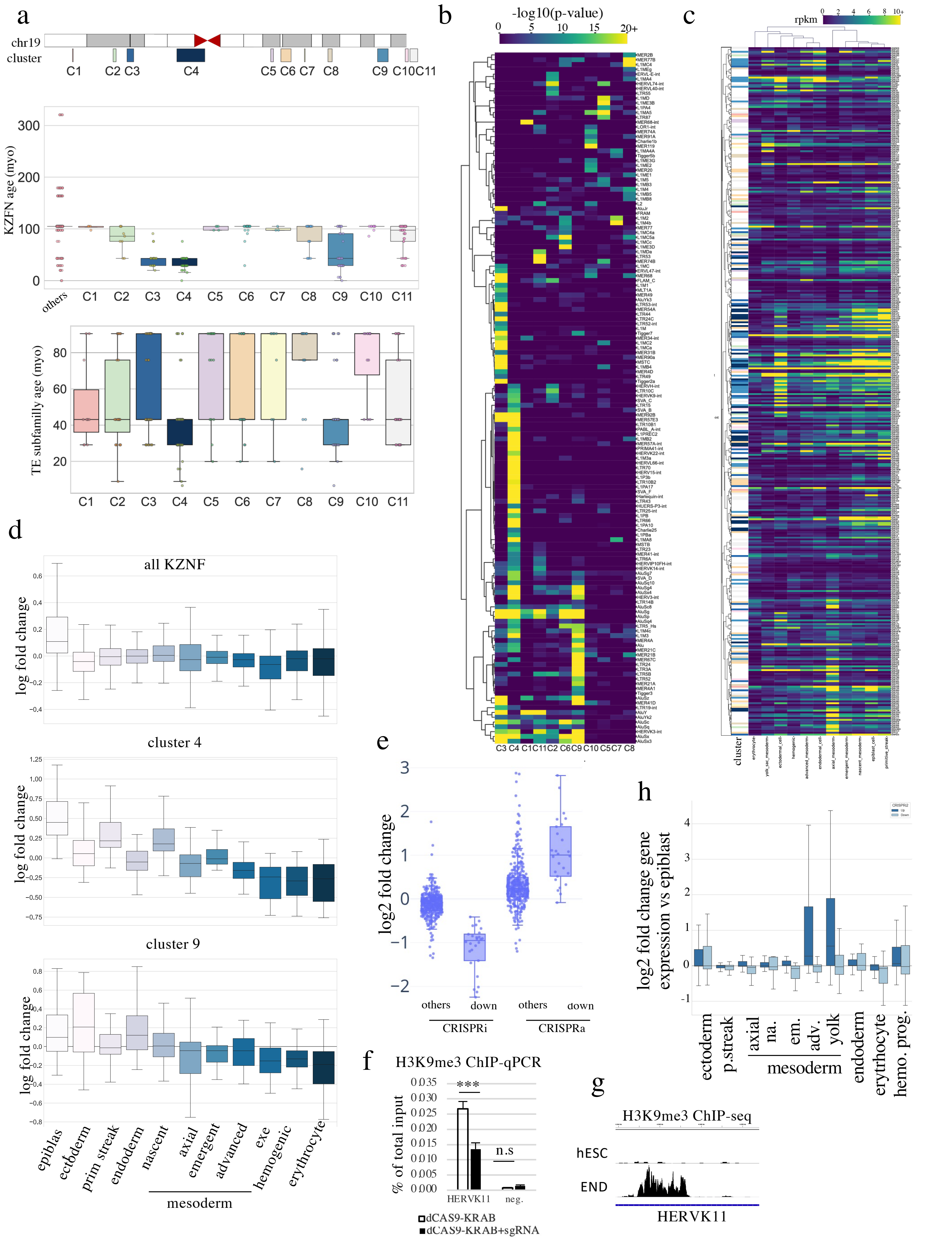
